# Sexually Dimorphic Response to Dietary Restriction-induced Longevity and Muscle Rejuvenation in *Nothobranchius furzeri*

**DOI:** 10.64898/2026.02.09.704879

**Authors:** Sonia Sandhi, Hannah Somers, Matthew Cox, Celeste Nobrega, Ryan Seaman, Elizabeth Bakers, Olivia Letchner, Robyn Reeve, Romain Menard, James Godwin, Anastasia Paulmann, Aric Rogers, Dario Riccardo Valenzano, Joel Graber, Hermann Haller, Romain Madelaine

## Abstract

Age-related skeletal muscle decline (sarcopenia) is a major contributor to frailty and mortality during aging, yet the extent to which sex shapes muscle aging and its response to dietary interventions remains poorly understood. Here, we use the short-lived vertebrate *Nothobranchius furzeri* (African turquoise killifish; ATK) to investigate how sex and intermittent fasting (IF) interact to regulate lifespan and skeletal-muscle aging. We establish and optimize an IF regimen that significantly extends lifespan in both male and female killifish, albeit with classical trade-offs including reduced growth and reproductive output. Despite these costs, IF markedly improves swimming performance in aged animals of both sexes. Structural analyses of killifish on a normal diet reveal pronounced sexual dimorphism in muscle aging. Males exhibit age-associated myofiber hyperplasia, whereas females maintain fiber number but undergo hypertrophic remodeling. IF partially reverses both phenotypes, restoring a more youthful fiber size distribution in both males and females. Single-nucleus RNA sequencing uncovers sex-specific remodeling of muscle-fiber composition in killifish on a normal diet, with females displaying an age-associated shift toward oxidative slow-twitch fibers that is reversed by IF, while males show relatively stable fiber-type proportions under normal and IF feeding regimens. Cell-cell communication analyses further reveal a global decline in intercellular signaling with age, alongside sex-specific restoration of distinct pathways under IF, including axon guidance and IGF signaling in females and metabolic ANGPTL signaling in males. Finally, bulk transcriptomic profiling demonstrates that aging follows largely sexually dimorphic molecular trajectories, whereas IF induces both sex-specific and shared responses. Notably, under IF, both sexes exhibit upregulation of ribosome biogenesis and genes supporting myofibrillar organization and contraction, likely underlying preserved muscle function. Together, these findings demonstrate that IF promotes longevity and muscle health through conserved anabolic mechanisms alongside sex-specific cellular and molecular rejuvenation strategies. Our work highlights the importance of incorporating sex as a biological variable when designing dietary interventions to promote healthy aging.

## INTRODUCTION

Age-related loss of muscle tissue, known as muscle atrophy, is a common hallmark of aging. The progressive decline of skeletal muscle mass and function, termed sarcopenia, is one of the most prominent features of aging (Cruz-Jentoft et al., 2019; Rosenberg, 1997). Sarcopenia manifests as reduced muscle-fiber size, impaired regenerative potential, and loss of neuromuscular interactions, leading to decreased mobility and increased risk of falls in the elderly population (Evans & Lexell, 1995; Larsson et al., 2019; Wilkinson et al., 2018). Beyond its impact on quality of life, age-related muscle atrophy contributes substantially to increased morbidity and mortality worldwide, highlighting its major clinical and social relevance (Brown et al., 2016; Sobestiansky et al., 2019). Despite decades of research, effective interventions to prevent or reverse muscle aging remain limited. Currently, regular physical exercise represents the most effective strategy to delay the onset and progression of sarcopenia, underscoring the importance of understanding the molecular and cellular mechanisms underlying muscle aging (Lavin et al., 2019; Mcleod et al., 2024).

Muscle aging does not proceed uniformly across sexes. In humans, men typically exhibit higher loss of fast-twitch fibers, while women show relatively more preserved integrity of the muscle tissue until menopause, after which muscle deterioration is accelerated by hormonal changes (Khadilkar, 2019; Nuzzo, 2024; Roberts et al., 2018; Rolland et al., 2007). These observations indicate a critical interplay between sex-specific biological factors—such as sexual hormones, genetic regulation, and metabolic differences—and the mechanisms underlying age-related muscle atrophy. Yet, the molecular bases of these sex-specific differences remain incompletely understood. Thus, understanding sex-specific trajectories impacting muscle homeostasis during aging is important for the development of tailored therapeutic approaches in human. However, the development of effective therapeutic strategies has been limited by the lack of short-lived vertebrate models to study the biology of muscle aging.

In recent years, the African turquoise killifish (ATK; *Nothobranchius furzeri*) has emerged as a powerful vertebrate model for aging research (Harel & Brunet, 2015; Reichard & Polačik, 2019; Valdesalici & Cellerino, 2003). Unlike traditional vertebrate models living many years or decades, *N. furzeri* has a remarkably short natural lifespan of only a few months. Despite the reduced lifespan, *N. furzeri* develop many hallmarks of aging observed in longer-lived species, including, but not limited to, increased age-related inflammation, cellular senescence, and skeletal-muscle atrophy (Hu & Brunet, 2018; Ma et al., 2025; Platzer & Englert, 2016; Ruparelia et al., 2024). This fish species also exhibits pronounced sexual dimorphism in muscle aging and recapitulates key features of muscle aging described in mammals, such as progressive muscle atrophy, depletion of the muscle stem-cell pool, and muscle-tissue denervation (Ruparelia et al., 2024). Together, these characteristics establish *N. furzeri* as an ideal system to study the biology of aging in a sex-specific context and investigate how sex-specific differences affect muscle aging, while providing translational relevance for human health.

Dietary restriction (DR) has emerged as a robust intervention to extend lifespan and delay age-associated functional decline in a wide range of invertebrate and vertebrate model organisms, ranging from *C. elegans* and *Drosophila* to mammals including rodents and primates (Colman et al., 2014; Kapahi et al., 2017; Lee et al., 2006; Swindell, 2012). Among the most extensively studied interventions capable of extending lifespan and healthspan are dietary strategies such as caloric restriction (CR) and intermittent fasting (IF). Beyond their effects on longevity, these dietary interventions also ameliorate age-associated pathologies, including neurodegeneration, metabolic dysfunction, and immune-system dysregulation (Martin et al., 2006; Zhang et al., 2024). In the context of skeletal-muscle aging, dietary-restriction strategies have been shown to preserve muscle-fiber size, improve mitochondrial function, and enhance autophagy(Chung & Chung, 2019; Gutiérrez-Casado et al., 2019; Lanza et al., 2012). However, despite increasing evidence of the benefits of dietary interventions, few studies have explored whether they exert sex-specific effects on muscle aging.

A recent report in *N. furzeri* demonstrated that DR can significantly increase lifespan in males, highlighting the potential of dietary interventions as powerful modulators of vertebrate aging and longevity(McKay et al., 2022). However, this protocol has been only optimized for male ATK, overlooking potential sex-specific differences in response to dietary manipulations. Sex is a critical biological variable influencing aging trajectories, metabolic regulation, and lifespan outcomes. For example, females exhibit distinct metabolic and hormonal regulation that may alter their response to fasting interventions (Bazhan et al., 2019; Freire et al., 2020). Therefore it is essential to determine whether existing CR or IF protocols are effective in a sex-dependent manner. To fill this gap, we developed IF protocols that we tested in male and female ATK to uncover sex-dependent and -independent responses to DR that may lead to the development of strategies for promoting healthy aging in all humans.

To address the critical gap described above, we investigated the interplay between sex, diet, and muscle aging in *N. furzeri*. This study reports sex-specific differences in muscle aging and determines whether IF can mitigate age-related muscle decline in a sex-dependent manner. Our results indicate that IF extends lifespan in male and female *N. furzeri* and improves muscle function through both shared and sex-specific mechanisms. Optimized DR enhanced longevity in males and females, while reducing growth and reproductive output. IF also improved swimming performance in both sexes, with males and females displaying distinct patterns of muscle remodeling involving hyperplasia and hypertrophy that can be reversed by IF. Single-nucleus RNA-sequencing and cell-cell communication analyses revealed sex-specific changes of myonuclei composition and signaling pathways with age and under DR. Transcriptomic profiling confirmed sexually dimorphic aging trajectories, yet both sexes shared upregulation of ribosomal activity and genes supporting muscle contraction after IF. Overall, IF promotes lifespan extension and muscle health through a combination of sex-specific and conserved cellular and molecular responses, underscoring the importance of considering sex in dietary interventions targeting aging to increase healthspan.

## RESULTS

### Optimization of intermittent-fasting induced longevity in males and females

We established intermittent fasting (IF) protocols that could potentially promote longevity in both male and female ATK. Adult males were housed individually (3-liter tanks), while females were housed in groups of 3 (6-liter tanks). Males and females were mated once every week for ∼12 hours. Between 30 and 40 days post-hatching (dph), all fish were acclimatised to an every-other-day feeding protocol before being sorted into distinct dietary regimes: IF1 and IF2. The first dietary intervention (IF1) consisted of two consecutive days of fasting followed by one day of feeding. The second dietary intervention (IF2) involved four days of fasting followed by three days of feeding. Control fish were maintained under *ad libitum* (AL) conditions, receiving daily feeding without prior acclimatisation period (Fig. 1A).

**Figure 1.**
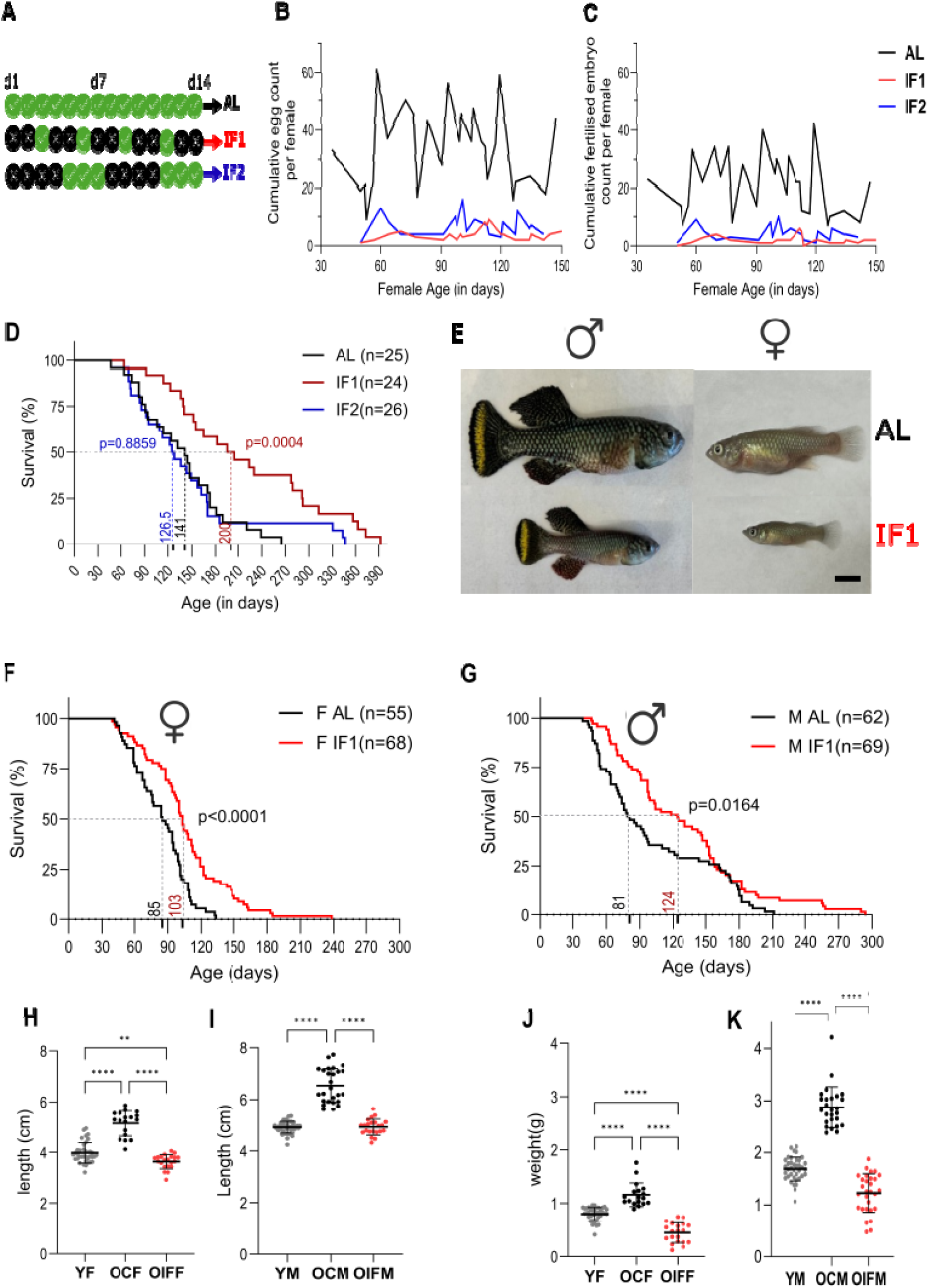
Optimisation of intermittent fasting method and trade-offs of longevity. **A)** Three different dietary regimes: AL - *ad libitum*, IF1 - 2 days fasting, 1 day feeding, IF2 - 4 days fasting, 4 days feeding. **B)** Average egg count per female and **C)** Average of fertilized embryo count per female on the Al, IF1, and IF2 regimes, from 30 dph to 150 dph. **D)** Kaplan Meier curve of combined male and female cohorts on the AL (n=25), IF1 (n=24), and IF2 (n=26) regimes with median survival at 141, 200, and 126.5 dph, respectively. T-test comparison of median lifespan of IF1 vs. AL and IF2 vs. AL, with p significance of 0.0004 and 0.8859, respectively. **E)** Representative images of male and female killifish at 90 dph under the AL and IF1 regimes. **F)** Kaplan Meier curve of females on the AL and IF1 regimes, with n=55 and 68, respectively. T-test for median lifespan with p<0.0001. **G)** Kaplan Meier curve of males on the AL and IF1 regimes, with n=62 and 69, respectively. T-test for median lifespan with p=0.0164. **H, I)** Total body-length measurements and **J), K)** body-weight measurements of females and males, respectively, at three different time points: young, old, and old IF1. One-way ANOVA with Tukey’s multiple comparison between samples, p value significance on bars represents *P ≤ 0.05, **P ≤ 0.005, ***P ≤ 0.0005.

We first assessed the effects of these different feeding regimes on reproductive capacity, by analyzing fertility from the onset of sexual maturity (40 dph) until advanced age (150 dph). Regime-matched fish of each sex were paired once per week to quantify reproductive performance by recording both the number of eggs laid and the proportion of eggs successfully fertilized across the lifespan of the fish. Both IF1 and IF2 protocols resulted in a decline in the cumulative number of eggs laid by females as well as in the number of fertilized embryos (Fig. 1B and 1C). Next, we analyzed the impact of the regimes on longevity of the fish, grouping males and females together in the analysis. We determined that the IF1 protocol significantly increased fish longevity, with a 41.8% extension in median lifespan (Fig. 1D). In contrast, the IF2 regimen reduced median lifespan by 10.3% (Fig. 1D). This observation highlights the importance of optimizing IF protocols for beneficial effects on longevity. Altogether, these results indicate that IF1 promotes lifespan extension at the expense of reproductive fitness, demonstrating the classical trade-off between reproduction and longevity.

Based on the results above, we selected IF1 as our method to promote longevity in *N. furzeri*. We repeated the experiment with larger cohorts and separated males and females for the analysis. We observed a significant increase in lifespan in both males and females, with a median lifespan extension of 53.08% in males and 21.18% in females (Fig. 1E–G). Next, we measured body length and body weight in young female (YF) and young male (YM) fish at 45 ± 2 dph and compared the values with *ad libitum*–fed controls (OCF, OCM) and old IF-fed fish (OIFF, OIFM). We found that while body length and weight naturally increased with age in both sexes, IF treatment led to a reduction in growth and weight in both sexes, with male body length and weight decreasing by 24.77% and 57.21%, respectively, and female body length and weight decreasing by 29.24% and 61.55%, respectively, compared to age-matched controls (Fig. 1H–K). These findings reveal a trade-off associated with IF-induced longevity, whereby lifespan extension is accompanied by reductions in both growth and reproductive capacity.

### Improved swimming performance of aged animals under IF feeding protocol

We next wanted to assess whether the IF protocol would improve muscle function. As a measure of muscle strength, we tested swimming performance using the swim tunnel assay (Fig. 2A and Fig. 2A’). In both male and female killifish, average swim speed declined with age but improved following IF treatment (Fig. 2B and C), by 60.86% and 41.15% in males and females, respectively. Remarkably, IF-treated old fish of both sexes swam faster and for longer periods of time than young controls. Across all age-matched groups, females outperformed males, indicating sex-specific differences in muscle performance in killifish during aging and DR.

**Figure 2.**
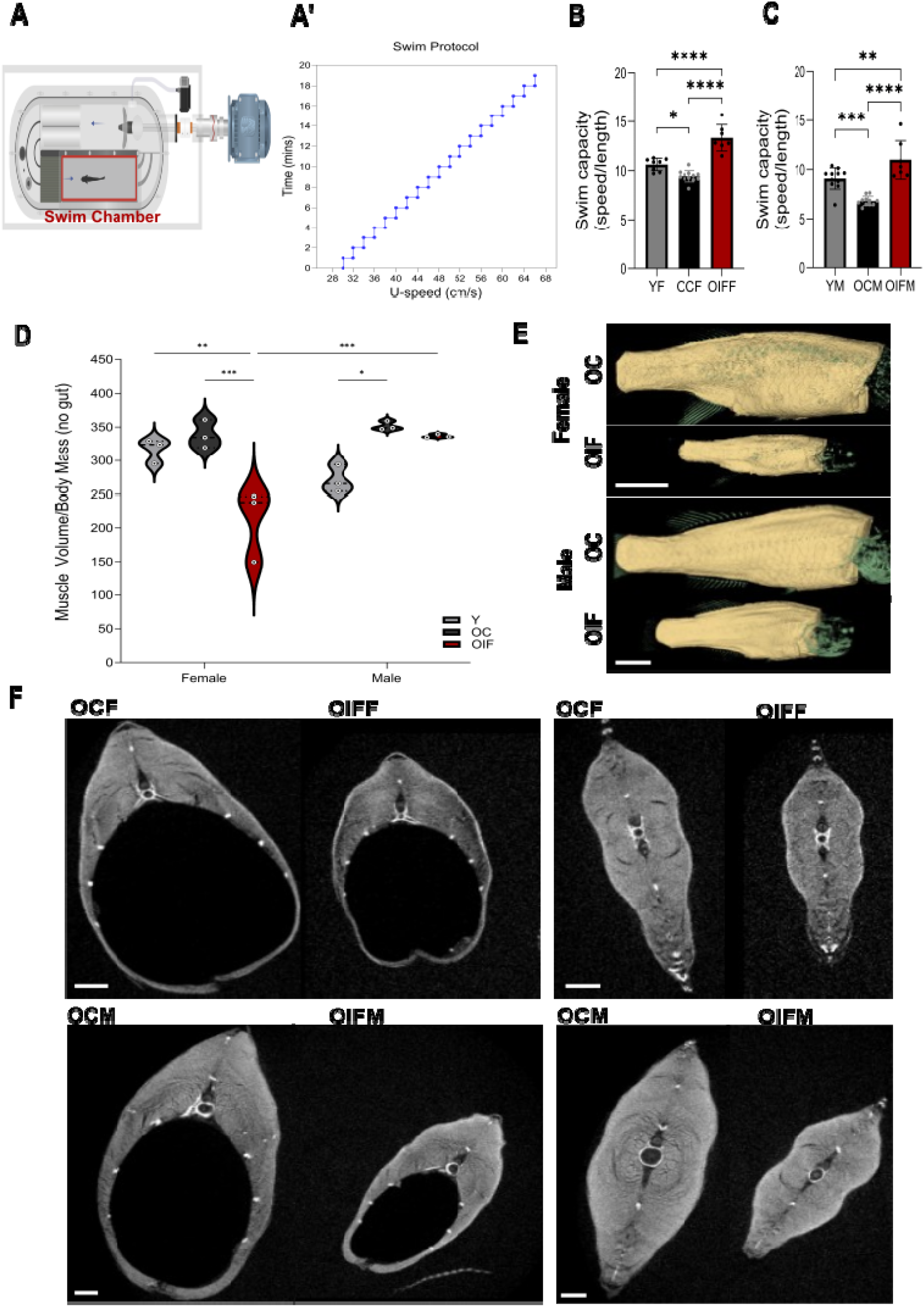
Improved swimming performance of aged animals under IF feeding protocol. **A)** Swim tunnel schematic (modified from Loligo systems) with arrows in the direction of water flow. **A’)** Swim protocol to check swim capacity of fish with water speed (cm/s) vs. time (in mins), with a step increase of 2cm/s every minute. **B)** Swim-speed to body-length ratio of females and **C)** males, at young, old control, and old IF treatments. Two-way ANOVA with Tukey’s multiple comparison between samples. **D)** Muscle-volume (mm^3^) to body-mass (g) ratio after removing internal organs and eggs (when present). Two-way ANOVA (treatment vs. sex) with Tukey’s multiple comparison within sex (F(2, 12) = 11.76, p = 0.0015). **E)** Representative 3D models of muscle measured. Muscle: yellow. Bone: green. Scale bar = 5 mm. **F)** Representative µCT slices of body-wall musculature (directly in front of the pelvic fin) and tail-base muscle (through the second caudal vertebra behind body cavity). Scale bar = 1 mm. p value significance on bars represents *P ≤ 0.05, **P ≤ 0.005, ***P ≤ 0.0005.

The skeletal muscle index (SMI)—defined as the ratio of muscle volume to body mass—is an indicator of favorable body composition and is generally associated with improved health and physical performance in humans and other species (Cruz-Jentoft et al., 2019; Wang et al., 2020). We therefore hypothesized that the enhanced swimming performance in IF-treated animals might result from a higher SMI. To test this, we performed microCT scans on the muscle tissue and observed that SMI remained unchanged between young and old control females, whereas it increased with age in control males (Fig. 2D-F), indicating a sex-specific response during aging. Under IF treatment, SMI decreased in females but remained unchanged in males (Fig. 2D-F), revealing another difference in the response to IF between males and females. These findings reveal interesting sex-specific differences in the maintenance of muscle mass with age and IF treatment in killifish and indicate that a higher mass of muscle tissue did not always correlate with improved swimming performance.

### IF feeding protocol reverses dimorphic muscle hyperplasia and hypertrophy

Because differences in SMI do not fully correspond to differences in swimming capacity, we next examined whether the number of muscle fibers declines with age, a phenomenon that could contribute to the age-associated reduction in swimming performance, and whether IF preserves myofiber number in aged animals.

To quantify myofibers, we stained cross sections from the caudal-muscle fin region using phalloidin-555 and WGA-647 (Fig. 3A). In female killifish, the total number of muscle fibers per myotome remained unchanged across ages and IF treatment conditions (Fig. 3B). In contrast, male killifish showed an age-dependent increase in total myofiber number (Fig. 3B), suggesting hyperplastic growth with aging that can be reversed by IF. Interestingly, both males and females exhibited an increase in large myofibers during aging (Fig. 3C, D). This hypertrophic phenotype can be alleviated by the IF feeding protocol in both males and females (Fig. C, D). We also determined that only males show an increase in the number of small fibers on a normal diet (Fig. 3C), supporting the male-specific hyperplasia phenotype mentioned above. Together, these data suggest that male and female killifish use different cellular mechanisms for muscle growth during aging, but that both muscle hyperplasia and hypertrophy can partially be reversed by DR.

**Figure 3.**
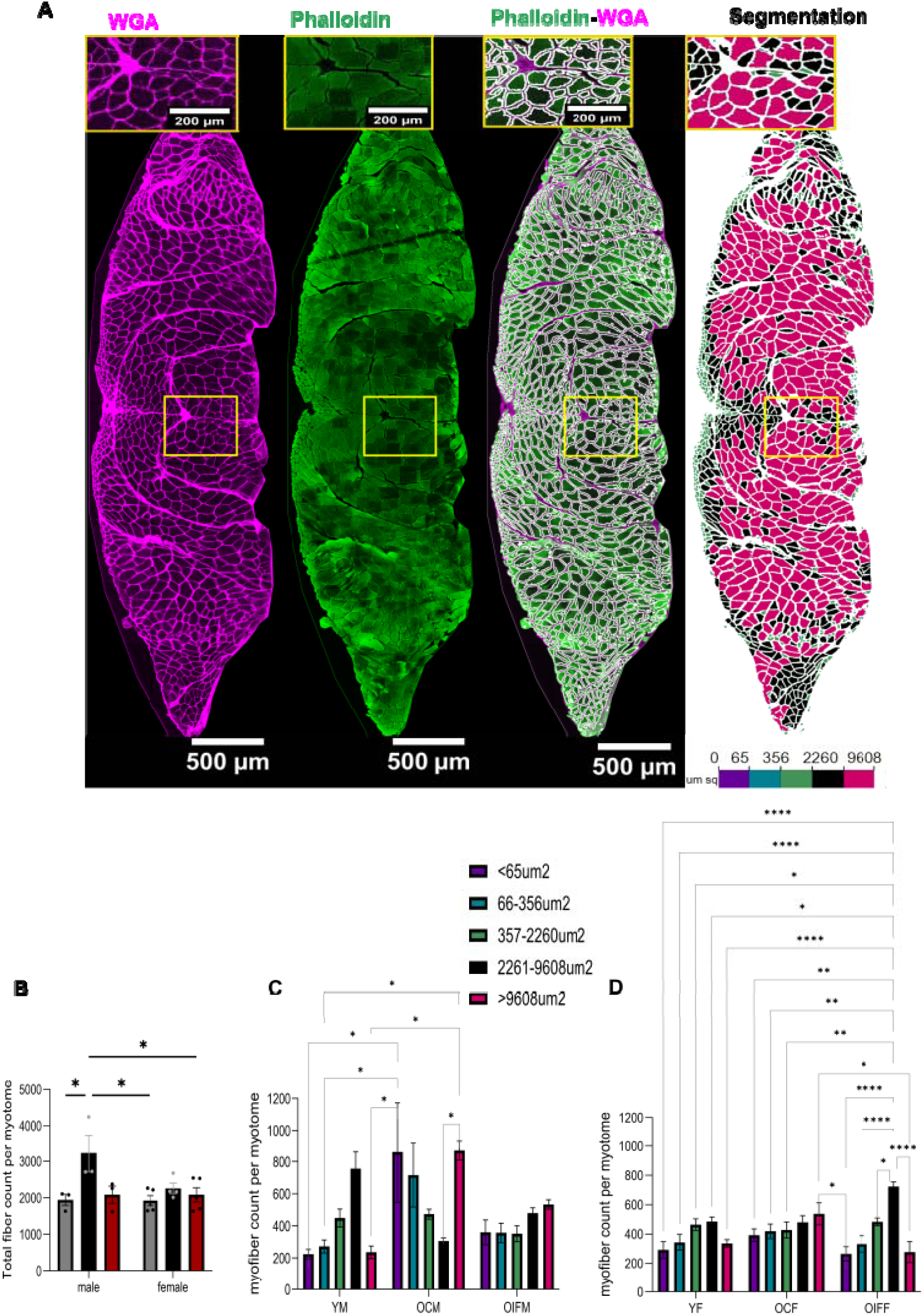
IF feeding protocol reverses sexually dimorphic muscle hyperplasia and hypertrophy. **A)** Representative images of myotome cross section stained with WGA, phalloidin, mask generated by subtraction of WGA from phalloidin and segmentation mask with color-coded fibers by size. **B)** Comparison of total fiber count per myotome between males and females, and across young, old, and old IF samples. **C, D)** Comparison of 5 different myofiber sizes: less than 65um sq., 66-356 um sq., 357-2,260 um sq., 2,261-9,608 um sq., and greater than 9,681 um sq., between young, old, and old IF samples in males (C) and (D) females. Two-way ANOVA, with Tukey’s multiple comparison, p value significance on bars represents *P ≤ 0.05, **P ≤ 0.005, ***P ≤ 0.0005.

### Sex-specific muscle-fiber-type remodelling by IF in aged animals

To characterize the cellular composition of killifish skeletal muscle, we performed single-nucleus RNA sequencing (snRNA-seq). Clusters were visualized using UMAP and annotated based on marker genes, identifying 11 different cell populations in killifish muscle (Fig. 4A and Fig. S1A-C). Across all conditions, including sex, age, and IF treatment, we did not detect any unique cell types specific to a particular group of animals, i.e., young control, old control, and old IF killifish (Fig. 4C,E), indicating that overall cell-type diversity is largely conserved in males and females, both with and without IF treatment. This analysis will provide a valuable resource of information for the scientific community using the ATK as a model of aging.

**Figure 4.**
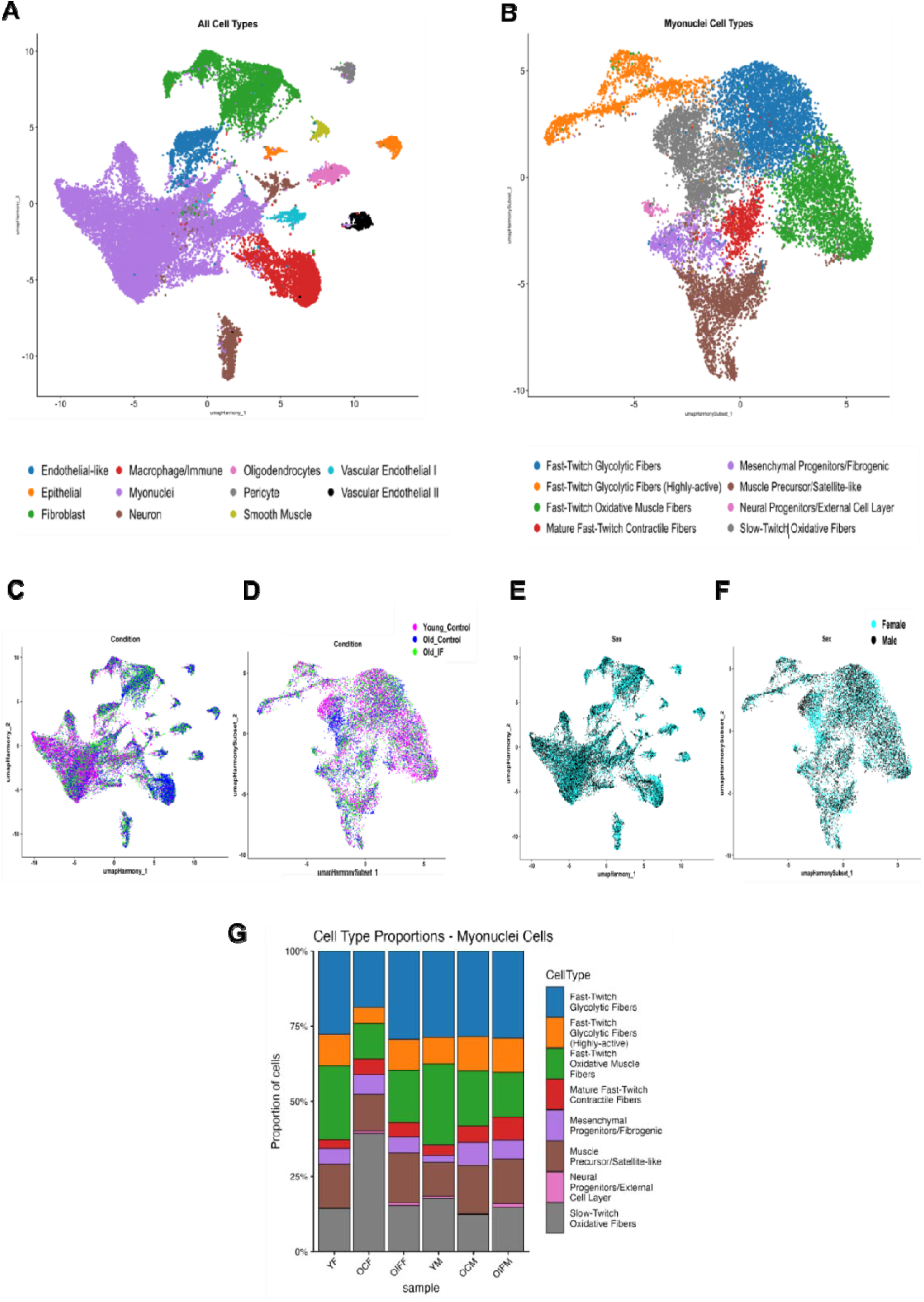
Myofiber composition is affected during aging and partially rejuvenated after IF. **A)** UMAP projection of single-nucleus RNA-sequencing data from young, old, and old IF, male and female killifish muscle reveals 11 distinct cell clusters, annotated by cell type. **B)** UMAP projection of single-nucleus RNA-sequencing data from young, old, and old IF, male and female killifish myonuclei subset reveals 8 distinct cell clusters. **C, E)** UMAP plot of all cell types and, **D, F)** only myonuclei subset, colored by regime condition, young, old, and old IF, or sex respectively, shows overlapping regions and differences between male and female transcriptomes. **G)** Stacked bar plots showing proportions of myonuclei-subset cell types across young, old, and old IF, males and females.

We next focused on myonuclei and performed sub-clustering to resolve distinct muscle-fiber types (Fig. 4B, D, and F, and Fig. S1D). We showed that muscle-fiber proportions decline with age in both male and female killifish, and that this effect is partially reversed by IF treatment (Fig. 4G). Analysis of fiber-type proportions revealed sex-specific and age-related patterns. In female control fish, both fast-twitch glycolytic and fast-twitch oxidative fibers declined with age, whereas slow-twitch oxidative fibers increased (Fig. 4G), suggesting an age-associated shift toward more oxidative fiber types in females. These modifications may represent a compensatory shift toward more fatigue-resistant, oxidative metabolism as muscle capacity declines with age. Importantly, IF treatment largely reversed these age-related changes in females (Fig. 4G), restoring a higher proportion of fast-twitch fibers at the expense of slow-twitch fibers. In contrast, male muscle displayed a different trajectory, as the proportion of the different muscle fibers was largely unaffected by aging and IF (Fig. 4G). Altogether, these results demonstrate that muscle-fiber composition is impacted differently in male and female killifish during aging.

Moreover, IF also exerts sex-specific effects, promoting fiber-type remodeling in females but not in males. These results highlight the importance of considering sex as a biological variable when assessing interventions aimed at mitigating age-related muscle decline.

### Sexually dimorphic intracellular-communication response after IF treatment

To explore how intercellular communication within muscle tissue changes with age and IF, we performed CellChat analysis (Jin et al., 2021) on the single-nuclei RNA-seq data. This approach allowed us to determine both the number and strength of cell-to-cell signalling interactions. We determined that the overall number of interactions declined with age in both male and female control killifish (Fig. 5A, B and Fig. S2A, B). However, we also identified sex-specific differences. The interaction strength was reduced in both females and males, but to a larger extent in females (Fig. S2A’, B’). Surprisingly, neither the number nor the strength of interactions was substantially reversed by IF (Fig. 5A, B and Fig. S2A’, B’). Further pathway-level analysis of cell-cell communication networks in the muscle tissue revealed that aging led to reduced information flow from different signalling pathways in males and females (Fig. 5C, D, and Fig. S2C). Interestingly, IF treatment reactivated several of these pathways in aged animals, but here again, males and females show differences in the pathways most strongly regulated by the feeding protocol. For example, signaling molecules involved in axon guidance such as netrin, ephrin, and semaphorin are downregulated in aged females and reversed by IF, but these molecules are neither downregulated with age nor reversed by IF in aged males (Fig. 5C, D). Remarkably, these signaling cues are received primarily by fast-twitch fibers, suggesting that functional interaction between neurons and muscle fibers may be disrupted during aging. Signaling dependent on growth hormones such as VEGF and EGF also declines with age in both males and females (Fig. 5C, D). Surprisingly, IF treatment reversed this effect only in females, indicating a sexually dimorphic tissue-growth response. We also showed that IGF pathway activity returned to a young-like state after IF treatment in old females, and that this was the case for ANGPTL pathway activity in males (Fig. 5C, D). Both the IGF and ANGPTL pathways regulate lipid and glucose metabolism(Choi et al., 2025; Xu et al., 2005), indicating that regulation of these processes may be a key component underlying restoration of the homeostatic condition in muscle tissue. Together, this analysis suggests that enhanced cellular crosstalk may contribute to the maintenance of muscle integrity and function during aging, and that males and females use different strategies to achieve the shared goal of muscle rejuvenation by regulating anabolic and catabolic processes.

**Figure 5.**
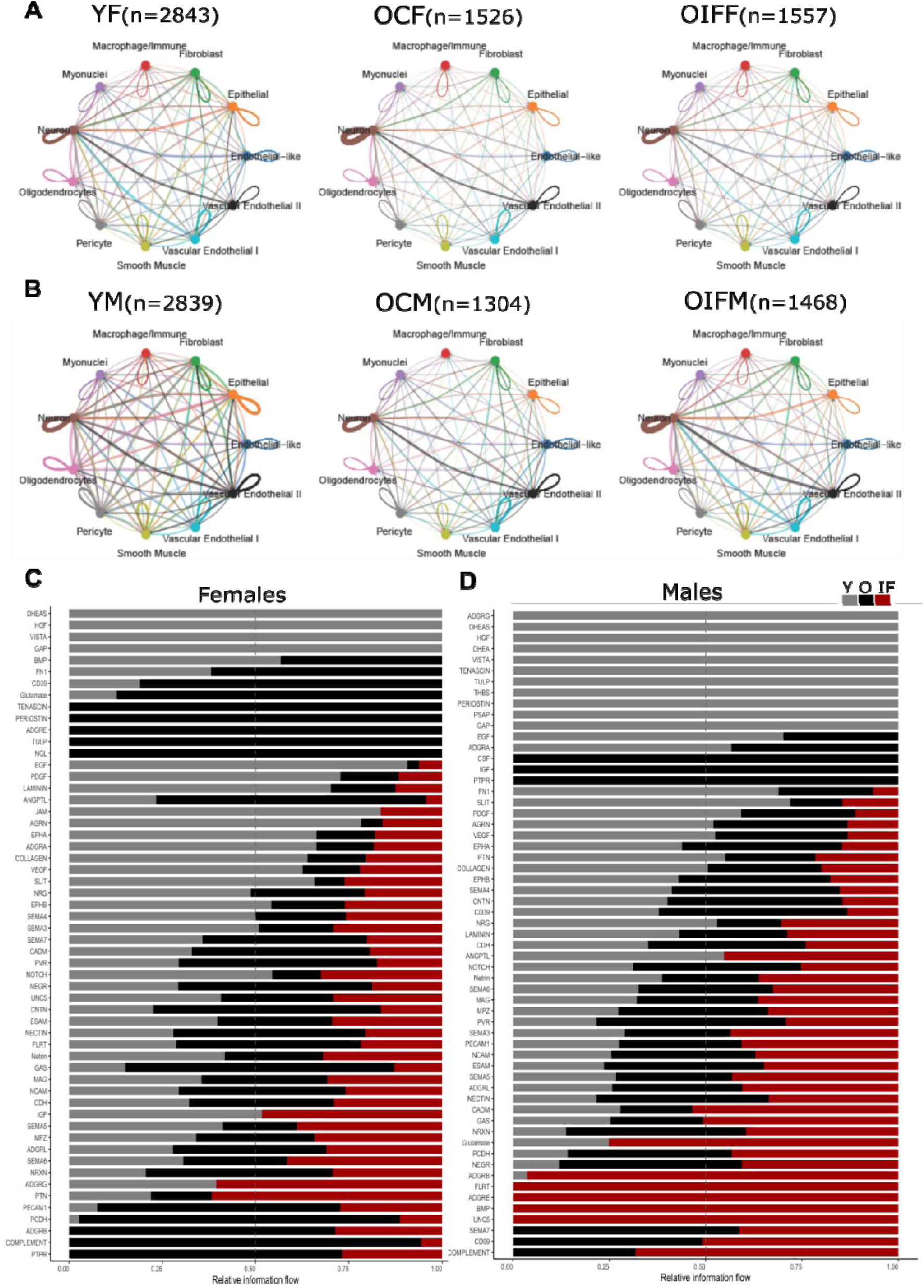
Cell-cell interactions decline with age and are improved by IF treatment. **A, B)** Circle plots visualize intercellular communication networks in young, old, and old IF killifish, in females and males, respectively, with the total number of interactions (N values) at the top of each circle plot. Each dot represents a cell type, with each connecting line representing a significant interaction between them. **C, D)** Bar graphs show relative information flow of individual cell-cell signaling pathways in young (grey), old (black), and old IF (red) in muscle cell types, of females and males, respectively.

### Sexually dimorphic transcriptional responses during aging and after dietary restriction

We next sought to investigate changes in gene-expression profiles underlying the sexually dimorphic response that we observed after the IF feeding protocol. To look at gene-expression changes with age, sex, and IF treatment, we performed bulk RNA sequencing using muscle tissue and compared transcriptomic profiles between males and females under different conditions (aging, feeding regimens). We identified 339 genes that were significantly dysregulated during aging in males (197 up and 142 down), and only 88 genes that were significantly dysregulated in females (32 up and 55 down) (Fig. S3A, B and Table S1-4). Among these genes, four collagen genes were downregulated in males and six in females, with *col1a1b*, *col1a2*, and *col2a1* being downregulated in both sexes, indicating that extracellular matrix (ECM) composition is affected in both males and females during aging. When comparing genes dysregulated in males and females during aging (Fig. 6E and Table S5, 6), we observed that only seven genes (including *col1a1b*, *col1a2*, and *col2a1*) are upregulated in both males and females and only two genes are downregulated in both sexes. Supporting this observation, our GSEA analysis indicated that killifish exhibit a sexually dimorphic response to biological processes affected strongly during aging, with rRNA processing being the most strongly upregulated process in females (Fig. 6A, B). This analysis demonstrated that many dysregulated processes are not following the same trajectory in males and females. Altogether, this transcriptomic analysis indicates that while ECM remodelling is a common feature of aging in male and female killifish, the aging trajectory differs between males and females.

**Figure 6.**
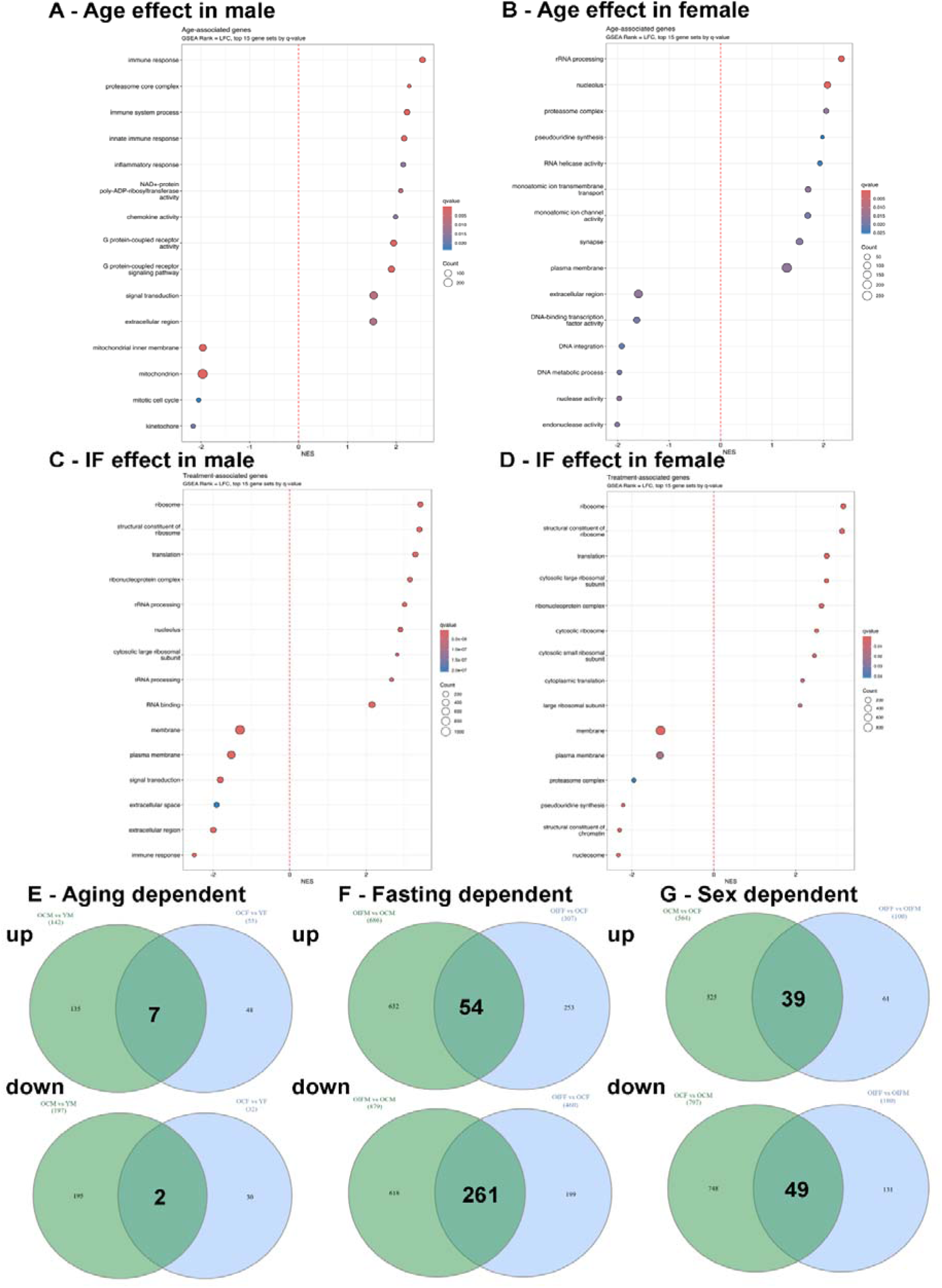
Sexually dimorphic changes in gene expression during aging and after if. **A, B)** GO (Gene Ontology) identified by GSEA analysis between young and old males (A) and females (B). **C, D)** GO (Gene Ontology) identified by GSEA analysis between old and old-after-IF males (C) and females (D).**E,F,G)** Graphical representation using Venn diagrams of the intersection in gene expression dependent on aging (E), fasting (F), and sex (G).

To assess similarities and differences between old males and females in their response to IF, we performed differential gene-expression analysis (Fig. 6F). We identified a total of 1,573 genes dysregulated in males (881 up and 692 down) and 769 in females (462 up and 307 down) in response to IF (Fig. S3C, D and Table S7-10). Interestingly, two of the top upregulated genes in both males and females after IF are *smyd1a* and *smyd1b,* two histone methyltransferases (Table S11, 12), which are critical regulators of myofibril organization and muscle contraction(Cai et al., 2019; Paone et al., 2018; Tan et al., 2006), potentially underlying improved swimming capacities after IF in both sexes. Another gene involved in muscle contraction, *mylpfb*, is upregulated in both males and females after IF (Table S11, 12); this gene is required for myofibril growth and fast-twitch function(Chong et al., 2020). Surprisingly, canonical target genes of the glucocorticoid-dependent stress-response pathway, *klf9* and *fkbp5*, are regulated in the opposite manner after IF treatment in old animals, indicating that, after IF, the stress response is downregulated in males while it is upregulated in females. Interestingly, we observed that IF reduced the differences in the gene expression profiles between males and females. When assessing the impact of aging and IF on gene expression in males and females, grouped together, we identified 1,366 genes dysregulated in young vs. control animals and only 285 genes dysregulated after IF (Table S13-16), indicating that the DR-induced longevity correlates with reduced heterogeneity in gene-expression differences between old males and females. Among these genes, only 88 are dysregulated in the same direction, i.e., either up or down, in both males and females, including fish under control conditions and those under IF (Fig. 6G), suggesting that these genes reflect differences between males and females that are not affected by the DR protocol. While females show enrichment for the ribosome biogenesis genes *rps* and *rpl* compared to males, both sexes show upregulation of genes associated with ribosome biogenesis after DR (Table S17, 18). Comparison of genes dysregulated in old males and females after IF identified 54 genes upregulated in both males and females and 261 genes downregulated in both sexes (Fig. 6F and Table S11, 12), highlighting the sexually dimorphic response in regulation of gene expression after fasting. Our GSEA analysis revealed that many of the top processes affected by IF in males and females are associated with ribosome biogenesis and functions such as translation (Fig. 6C, D). A significant difference between males and females in the response to IF is that terms associated with cell communication and ECM, such as signal transduction and extracellular space, are downregulated in males but not in females. Altogether, these data indicate that while males and females exhibit many differences in their responses to the same fasting protocol, upregulation of ribosomal activity is a likely a shared phenomenon underlying increased muscle health and function after DR.

This work sheds light on how sex and diet interact to shape the progression of muscle aging in a short-lived vertebrate. Beyond advancing our fundamental understanding of muscle biology, these findings have translational implications for the design of sex-specific interventions aimed at preserving muscle health and promoting healthy aging in humans.

## DISCUSSION

Aging of skeletal muscle is characterized by progressive loss of muscle mass, function, and cellular homeostasis, yet accumulating evidence indicates that these trajectories differ substantially between sexes. Using the short-lived vertebrate *Nothobranchius furzeri*, we systematically examined how sex and intermittent fasting (IF) interact to shape muscle aging. Our multi-scale analysis—integrating physiology, histology, and bulk and single-nucleus transcriptomics—reveals that males and females follow distinct structural and molecular aging pathways and respond to dietary restriction through partly shared but predominantly sex-specific mechanisms. These findings position *N. furzeri* as a relevant model for exploring sexually dimorphic determinants of muscle aging and highlight the need for sex-aware intervention strategies.

### Sex-specific trajectories of muscle aging

We show that male and female muscles follow fundamentally different pathways of structural remodeling during aging. Males exhibit a marked hyperplastic response, increasing myofiber number with age, whereas females preserve fiber number but display hypertrophy of existing fibers. Both sexes accumulate large fibers with age, consistent with age-associated impairments in neuromuscular connectivity and regenerative capacity documented in other vertebrate systems (Aare et al., 2016; Kelly et al., 2018). Our transcriptomic profiling further reveals that males undergo substantially broader gene-expression remodeling during aging than females.

Although both sexes show downregulation of several ECM components—a shared hallmark of aged muscle—the majority of age-associated changes are sex-specific. Processes such as rRNA metabolism follow opposing trajectories in males and females, supporting the notion that muscle aging in *N. furzeri* is not merely quantitatively different between sexes but rather is governed by distinct sexually dimorphic molecular programs.

### IF extends lifespan but imposes canonical life-history trade-offs

Optimizing fasting schedules allowed us to identify an IF protocol that extends lifespan in both sexes but reduces growth and reproductive output, consistent with classical trade-offs between somatic maintenance and reproductive investment.

Despite reduced body size, IF-treated fish of both sexes showed improved swimming performance at old age. Importantly, these functional gains occurred independently of skeletal muscle index, suggesting a qualitative rejuvenation of muscle tissue rather than simple preservation of mass. This dissociation between muscle mass and function parallels some human studies, where functional outcomes are more predictive of healthspan than muscle size alone (Newman et al., 2006).

### IF reverses structural hallmarks of aged muscle through sex-specific mechanisms

Our histological analyses demonstrate that IF normalizes age-associated hyperplasia and hypertrophy. In males, IF reduces the aberrant increase in small myofibers, suggesting a restoration of homeostatic control over fiber recruitment. In both sexes, IF alleviates hypertrophy, restoring a more youthful fiber-size distribution. Single-nucleus transcriptomics further reveals that IF induces a marked remodeling of fiber-type composition in females, reversing the age-associated shift toward slow oxidative fibers and restoring a youthful prevalence of fast-twitch glycolytic and oxidative fibers. In contrast, males show minimal changes in fiber-type proportions in response to IF. These findings suggest that improved muscle function in males is achieved primarily through structural and metabolic normalization, whereas in females, IF induces muscle fiber-type youthful remodeling.

### Cell-cell communication is remodeled in a sex-specific manner by aging and IF

Aging caused a broad reduction in intercellular signaling strength within muscle tissue; this decline was more pronounced in females. IF did not globally restore signaling but instead reactivated specific pathways in a sex-dependent manner. In females, IF re-engaged axon-guidance pathways—including netrin, ephrin, and semaphorin—primarily affecting fast-twitch fibers. This may reflect partial restoration of neuromuscular connectivity, consistent with the functional improvements observed. In males, IF selectively restored ANGPTL signaling associated with lipid and glucose metabolism, indicating metabolic recalibration as a potential rejuvenation mechanism. Activation of IGF signaling occurred predominantly in females, further underscoring distinct anabolic responses to fasting. Together, these pathway-level differences between males and females show that cellular communication networks deteriorate with age but can be reprogrammed by dietary intervention in a sex-specific manner.

### Sexually dimorphic transcriptional responses to IF reveal distinct rejuvenation strategies

Bulk RNA sequencing demonstrated widespread transcriptional reprogramming after IF in both sexes, with males showing nearly twice as many dysregulated genes as females. Despite this divergence, several features were shared: both sexes exhibited upregulation of genes involved in ribosome biogenesis and translation, as well as *smyd1a/b* and *mylpfb*, which promote myofibrillar organization and contractile function. These shared responses likely contribute to improved swimming capacity across sexes. However, ECM-associated pathways were selectively reduced in males, demonstrating that tissue remodeling in response to fasting is strongly sex-dependent. Additionally, stress-response pathways diverged notably: glucocorticoid-responsive genes were downregulated in males but upregulated in females, suggesting that fasting modulates stress signaling in opposite trajectories. Collectively, these transcriptomic analyses indicate that IF induces a common anabolic signature centered on enhanced ribosomal activity but otherwise elicits sex-specific molecular rejuvenation strategies—females primarily via neuromuscular and fiber-type remodeling, males via metabolic restructuring and suppression of catabolic signaling.

### Implications for modeling and treating muscle aging

Our findings establish *N. furzeri* as a powerful platform for studying sex-dependent muscle aging. The extensive dimorphisms observed across structural, functional, and molecular levels highlight important considerations for interventions targeting age-related muscle decline. Our finding that IF improves muscle function in both sexes but does so through mechanistically distinct pathways suggests that therapeutic strategies may require sex-tailored implementation to yield beneficial outcomes in humans. Moreover, the selective reactivation of neuromuscular signaling pathways in females and metabolic pathways in males underscores the need to understand how hormonal and intrinsic tissue factors govern these divergent responses.

## CONCLUSION

Overall, our findings are highly consistent with mammalian data demonstrating that dietary restriction extends lifespan, improves muscle function independently of mass, and engages conserved anabolic and metabolic pathways. Importantly, this study extends mammalian work by resolving sex-specific mechanisms at cellular and transcriptomic resolution, revealing that males and females achieve muscle rejuvenation through distinct yet convergent biological strategies. The conservation of fiber-type remodeling, neuromuscular signaling pathways, IGF and metabolic regulation, and ribosome biogenesis supports the translational relevance of *N. furzeri* as a vertebrate model for studying sex-specific interventions targeting sarcopenia and healthy aging.

**Table S1. Comparison between OCM and YM: genes down**

**Table S2. Comparison between OCM and YM: genes up**

**Table S3. Comparison between OCF and YF: genes down**

**Table S4. Comparison between OCF and YF: genes up**

**Table S5. Genes similarly downregulated during aging**

**Table S6. Genes similarly upregulated during aging**

**Table S7. Comparison between OIFM and OCM: genes down**

**Table S8. Comparison between OIFM and OCM: genes up**

**Table S9. Comparison between OIFF and OCF: genes down**

**Table S10. Comparison between OIFF and OCF: genes up**

**Table S11. Genes similarly downregulated during fasting**

**Table S12. Genes similarly upregulated during fasting**

**Table S13. Comparison between OCF and OCM: genes down**

**Table S14. Comparison between OCF and OCM: genes up**

**Table S15. Comparison between OIFF and OIFM: genes down**

**Table S16. Comparison between OIFF and OIFM: genes up**

**Table S17. Genes similarly downregulated in aged control and IF conditions**

**Table S18. Genes similarly upregulated in aged control and IF conditions**

## MATERIALS AND METHODS

### Animal husbandry and feeding protocols

All the work reported here was performed in the wild-type GRZ strain of African turquoise killifish (*Nothobranchius furzeri*). All procedures were approved by the MDIBL Institutional Animal Care and Use Committee (IACUC, #24-02) and comply with the MDI Biological Laboratory Institutional Assurance # D16-00341.

Fish were maintained at 27°C in an Aquaneering® recirculating system under a 12h light/dark cycle, and water parameters were maintained at pH 7.0–7.4, conductivity 2800 µS. Each cohort raised was randomly split at adulthood stage (30 dph) into different lifelong feeding regimes: AL, IF1 and/or IF2. Male fish after sexual maturity were single-housed in 3L tanks, while females were group-housed with n=2-3 in a 6L tank. Fish designated for any IF regime went through an acclimation period of 10-12 days with alternate days of feeding and fasting before entering the IF1 or IF2 regime. Male and female tanks were kept next to each other and fish were mated once each week, at the end of a feeding day.

Embryos were collected, disinfected in 550ppm PVP Iodine (Ovadine®, Syndel, USA), and incubated at 28°C in Ringer’s Solution. At the black-iris stage (12-14 days post fertilization (dpf)), embryos were transferred to coconut plates with Whatman® filter paper and stored at 27°C for three weeks. Golden-eyed embryos were hatched using a salt solution (1g Instant Ocean® in 1L RO water, pH=6, 4°C). Hatchings were raised offline for first week, and then on 7 dph (days-post hatching), they were transferred to the water system. Fish were fed Artemia and Otohime pellet food as per age-dependent protocols shown in Table 1. On feeding days, fish were fed 100uL of dense Artemia slurry and 3 pellet feedings of 20 mg per fish, with the last feeding at approximately 3:30pm. Any leftover food in the tanks was siphoned out at the end of the feeding day. On fasting days, no food was given to the fish.

**Table 1.**
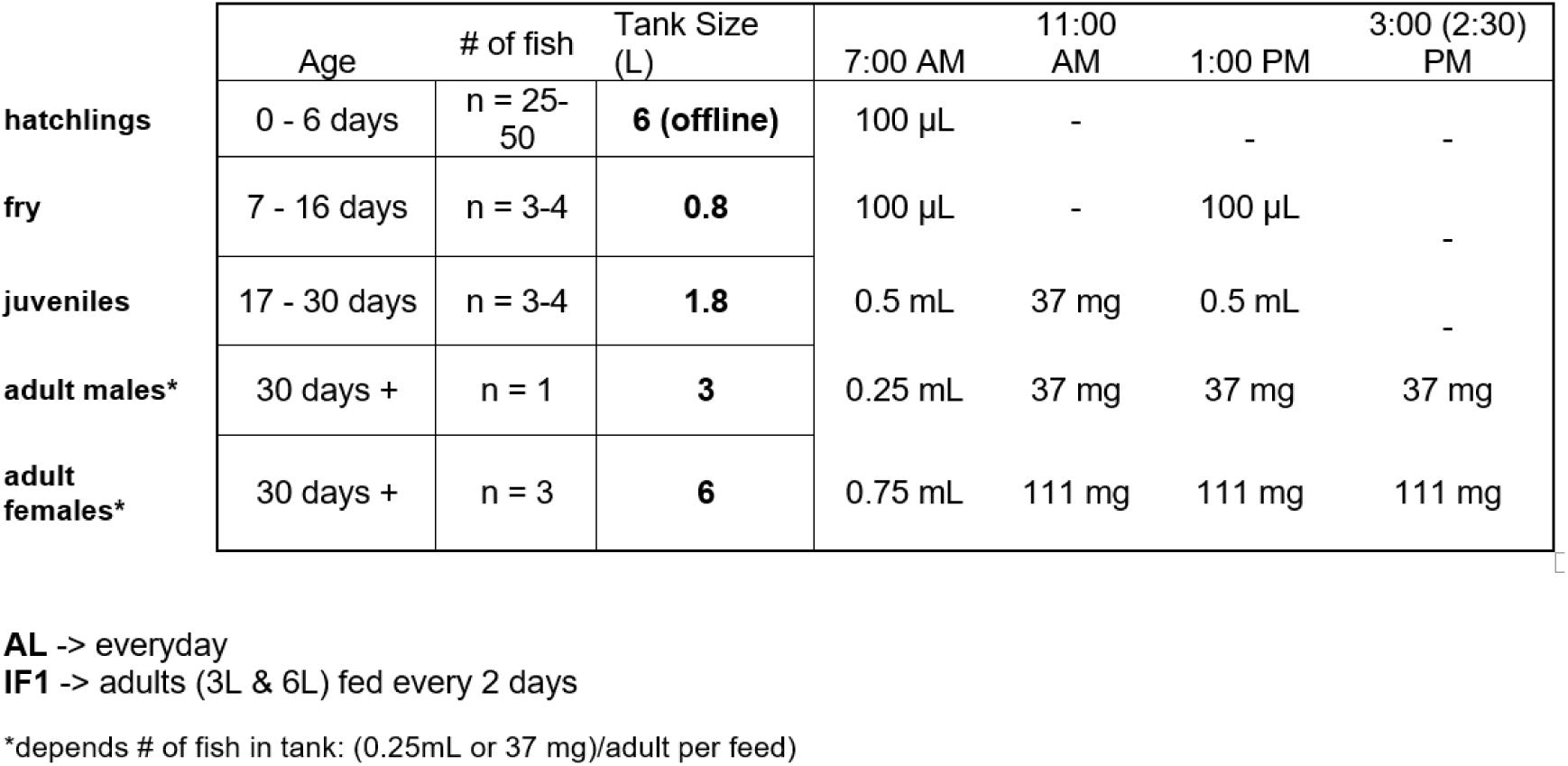
Feeding and housing schedule of different age groups of killifish. ATK feeding schedule indicating housing conditions and the quantity of concentrated artemia (in μl or ml) or Otohime C1 pellets (in mg) served for feeding.

|  | Age | # of fish | Tank Size (L) | 7:00 AM | 11:00 AM | 1:00 PM | 3:00 (2:30) PM |
| --- | --- | --- | --- | --- | --- | --- | --- |
| **hatchlings** | 0 - 6 days | n = 25-50 | **6 (offline)** | 100 µL | - | - | - |
| **fry** | 7 - 16 days | n = 3-4 | **0.8** | 100 µL | - | 100 µL | - |
| **juveniles** | 17 - 30 days | n = 3-4 | **1.8** | 0.5 mL | 37 mg | 0.5 mL | - |
| **adult males\*** | 30 days + | n = 1 | **3** | 0.25 mL | 37 mg | 37 mg | 37 mg |
| **adult females\*** | 30 days + | n = 3 | **6** | 0.75 mL | 111 mg | 111 mg | 111 mg |

**AL** -> everyday
**IF1** -> adults (3L & 6L) fed every 2 days
\*depends # of fish in tank: (0.25mL or 37 mg)/adult per feed)

### Length and weight measurements

Length and weight of fish were measured at young and old age, 45 do and 90 do, respectively. Fish were anesthetized using 0.2 g/L of MS-222 (Syncaine®, Syndal, USA) and weighed on an Adventurer™ Pro Analytical Electronic Balance (model: AV64C, Ohaus®, USA). Images for length measures were taken using a camera, with subsequent transfer of image data in .png format. Length measurements were performed using FiJi (v1.54)(Schindelin et al., 2012) from the tip of the snout to the end of the tail fin using the line tool.

### Histology

Fish were euthanized by 0.5 g/L of MS-222 (Syncaine®, Syndal, USA), followed by ice incubation for 10 minutes. Tail-fin muscle tissue was then cut from cloaca till the opposite end dorsal fin, fixed with 4% PFA solution in 1xPBS (Phosphate Buffer Saline) at 4°C overnight, and then incubated in 30% sucrose in PBS 1x solution at 4°C overnight, followed by OCT embedding and snap freezing using liquid nitrogen. −80°C stored samples were thin-sliced into 30-um cryosections and stained with WGA-647 (Invitrogen; W32466) and phalloidin-555 (Invitrogen; A34055) used at 1:100 and 1:300 dilution in PBS 1x solution, respectively.

### Imaging of myofibers

All images of the ATK muscle sections were collected with a Zeiss LSM 980 confocal microscope (Carl Zeiss Microscopy) on an upright Zeiss Axio Examiner stand equipped with a Zeiss Plan Apo 10x/0.45 NA objective (Carl Zeiss Microscopy).

WGA fluorescence was excited with the 561 nm line, 1% laser intensity from an 8-mW laser diode. Alexa 647 fluorescence was excited with a 639 nm line, 2% laser intensity from a 25-mW laser diode. Laser beam attenuation for all channels other were captured using Airyscan 2 with the following detection wavelengths: 561 from 422 to 477 nm and 573 to 627 nm, Alexa 647 from 499 to 557 nm and 659 to 720 nm.

Images were sequentially acquired in confocal mode at zoom 1.7, with a line average of 1, a resolution of 1024x1024 pixels, a pixel time of 0.77 µs, in 8-bit and in bidirectional mode. Z-stack images were collected with a step size that varied between images with the Motorized Scanning Stage 130x85 PIEZO (Carl Zeiss Microscopy) mounted on Z-piezo stage insert WSB 500 (Carl Zeiss Microscopy).

Tiles were collected based on the size of the muscle-fiber section. The microscope was controlled using Zen Blue software (Zen Pro 3.1), Airyscan images were processed automatically and saved in CZI file format.

### Image Analysis

Fiji (ImageJ v1.54f) was utilized for image preparation and analysis. Maximum intensity projections were created from z-stack acquisitions for all four channels. The polygon selection tool was used to manually outline each muscle section, and regions outside the area of interest were cleared. Rolling ball background subtraction (radius = 100 pixels) was applied to all channels. Images were then split into individual channels and saved as separate TIF files for downstream processing.

To improve segmentation quality, the phalloidin-555 and WGA-647 channels were normalized using Fiji’s Enhance Contrast function with 0.5% saturated pixels and both normalize and equalize options selected.

Ilastik v1.4.0.post1-gpu was used for supervised pixel classification segmentation. Two separate classifiers were trained: (1) five phalloidin images were annotated to identify muscle-fiber bodies, with the “suggest features” tool used to optimize feature selection; (2) five WGA images were annotated to identify fiber boundaries, also utilizing the “suggest features” tool for feature optimization. Both classifiers were applied via batch processing to all images, and simple segmentation probability maps were exported as tiff images.

Morphological refinement of the Ilastik segmentation masks was performed in Fiji using the MorphoLibJ plugin. The phalloidin mask was processed with morphological erosion (diamond element, radius = 1) followed by Gaussian blur (σ = 2) to enhance fiber regions. The WGA mask underwent morphological dilation (horizontal line element, radius = 1) to smooth fiber boundaries. The processed phalloidin mask was then subtracted from the processed WGA mask using the Image Calculator to isolate fiber boundaries. The resulting binary image was dilated (disk element, radius = 1) to close gaps between fiber segments, followed by binary hole filling to create complete fiber regions.

Individual muscle fibers were identified using the Analyze Particles tool with size constraints of 25-100,000,000 pixels² and minimum circularity of 0. Spatial calibration was set to 1.0 pixels/μm. Comprehensive morphometric measurements were collected for each fiber, including area, perimeter, circularity, minimum and maximum Feret diameter, centroid coordinates, bounding box dimensions, aspect ratio, roundness, and solidity.

Color-coded fiber size maps were generated using a percentile-based categorical scheme for visualization. Size category thresholds were defined from pooled analysis of fibers from old female control images: 20th percentile = 65 μm², 40th percentile = 356 μm², 60th percentile = 2260 μm², 80th percentile = 9688 μm². These thresholds were applied consistently across all experimental conditions. A colorblind-friendly categorical color palette (small: deep purple; medium-small: teal; medium: forest green; medium-large: black; large: deep magenta) was used for all visualizations.

All ROI sets and quantitative measurements were exported as CSV files for statistical analysis.

### Bulk RNA sequencing

Total RNA samples for bulk RNA-seq analysis were prepared with snap-frozen muscle tissue. Each experimental condition was performed using biological triplicates to ensure reproducibility. RNA extraction, library preparation, and sequencing were carried out by NovoGene using the Illumina NGS platform. Raw FASTQ files were processed using the nf-core/rnaseq pipeline (v3.19.0)(Ewels et al., 2020; Harshil Patel et al., 2025) with default parameters. Reads were aligned to the *Nothobranchius furzeri* reference genome (Nfu_20140520) obtained from Ensembl release 111. Prior to differential expression analysis, unwanted variation was removed using RUVseq (v1.42.0)(Risso et al., 2014). Differential gene-expression analysis was performed using DESeq2 (v1.48.2)(Love et al., 2014). Gene Set Enrichment Analysis (GSEA) and over-representation analysis (ORA) were conducted using clusterProfiler (v4.16.0)(Xu et al., 2024). All visualizations were generated using ggplot2 (v4.0.0)(Wickham, 2016) and enrichplot (v1.28.4)(Guangchuang Yu, 2018).

### Single nuclei RNA sequencing

Nuclei were isolated 165 mg each of flash-frozen muscle tissue collected from young and old, female and male fish put on either the control or IF feeding regime (n=3 fish pooled each sample), and we captured single nuclei and constructed 3’ single-cell gene-expression libraries (Next GEM-X v4) using the 10x Genomics Chromium system. The libraries were sequenced with ∼200 million PE150 reads on Illumina NovaSeq. The raw fastq-format sequencing data was processed, and the reads were aligned to the Ensembl (v111) African turquoise killifish reference using STAR implemented in the 10x Genomics Cell Ranger software.

R Studio was used to run custom R scripts for analysis. Genes expressed in fewer than three cells were excluded from the analysis. Cell quality control was conducted on the data by eliminating the bottom 10% of cells based on read count and gene count, along with cells containing a mitochondrial percentage greater than 10%.

Doublets were identified and removed using the *DoubletFinder* package (v2.0.4) (McGinnis et al., 2019). The *Seurat* package (v5.0.2)(Hao et al., 2021) was used for clustering and cell type identification. Expression levels were normalized through the “NormalizeData” function with the LogNormalize method. For visualization, the “ScaleData” function was used to regress out differences in the number of molecules, number of genes, percent mitochondrial genes, and cell cycle effects.

Principal component analysis (PCA) was conducted, and the first 19 PCs were used to identify clusters and generate both t-distributed stochastic neighbor embedding (t-SNE) and uniform manifold approximation and projection (UMAP) plots. UMAPs were displayed in Loupe Browser 8 (10x Chromium), and specific cell clusters were determined by the expression of marker genes that define specific cell types and known expression as known in zebrafish and other teleosts. Identified cell cluster were verified by using FindAllMarkers function with options min.pctc=c0.25, min.diff.pct=0.25.

The *CellChat* package (v2.1.2) (Jin et al., 2025) was used to infer intercellular communication within the single-nuclei data. The pre-curated human database for intercellular communication was translated into African turquoise killifish using orthology maps derived from Ensembl (v111). Non-one-to-one mappings were maintained in the translated database by duplicating interaction entries for each African turquoise killifish gene that was mapped to a Human gene.

Differentially expressed genes between cell types or samples were calculated using the “FindMarkers” function in *Seurat.* These genes were ranked in descending order by the negative base-ten logarithm of the *p*-value multiplied by the sign of their log_₂_ fold-change value. As no hallmark gene sets exist relating to *Nothobranchius furzeri,* the Gene Ontology (GO) terms from *Danio rerio* (Zebrafish) in the biological process, molecular function, and cellular component categories were translated to the killifish genome using orthology maps.

Gene set enrichment analysis was performed using the *clusterProfiler* package with the “gseGO” function. The minimum gene set size was set to ten and the maximum gene set size was set to 500. A *p*-value cutoff of 0.05 was applied to identify significantly enriched gene sets. Because redundant gene ontology terms were common, these were reduced using the *rrvgo* package (v 1.21.5) (Sayols, 2023). A similarity matrix of GO terms was first calculated using the “calculateSimMatrix” function, and redundant terms were then summarized with the “reduceSimMatrix” function. Plots were generated using the *ggplot2* package(Wickham, 2016).

### Swim test

Swim tunnel (Loligo system-SW10100) was used to look at the swimming performance of young fish and old fish, under control feeding and IF treatment. The system calibration and maintenance were adapted from *Burris et al*(Burris et al., 2021). The swimming performance assay was modified to use the velocity increments of 2cm/s every 1 minute, with the use of 3cm length, to record the maximum swim speed reached by each individual fish. Each fish was acclimated in the swim chamber for 15 minutes before the start of the experiment with no flow. Fish which did not swim were removed from the experiment.

### microCT scan

Animals were scanned on a Bruker Skyscan 1272 micro-CT using an aluminum 0.5mm filter with 0.2° rotation step, random movement of 30, and frame averaging of 4. Automatic thresholding was used to remove bone and background from the body muscle mass. The head and associated non-muscular tissue were removed by cropping behind the gill arch and in front of the second rib. All fins and associated muscles outside the body wall were removed and any soft tissue or eggs within the body wall that were left after gutting were also manually removed. Models and muscle volume were generated using 3D Slicer 5.8.1.

## CONTRIBUTIONS

*Conceptualization*: Sonia Sandhi, Aric Rogers, Dario Valenzano, Romain Madelaine

*Investigation*: Sonia Sandhi

*Data Collection*: Sonia Sandhi, Elizabeth Bakers, Olivia Letchner, Hannah Somers, Robyn Reeve, James Godwin, Romain Menard, Anastasia Paulmann

*Data Analysis*: Sonia Sandhi, Celeste Nobrega, Matthew Cox, Ryan Seaman, Joel Graber, Hannah Somers

*Writing*: Sonia Sandhi, Romain Madelaine

*Funding acquisition*: Aric Rogers, Hermann Haller, Romain Madelaine

*Supervision*: Aric Rogers, Hermann Haller, Romain Madelaine

## DATA AVAILABILITY STATEMENT

Data availability. All data presented in this manuscript are available to the scientific community, either in a public repository, within the manuscript itself or as supplementary information. The transcriptomics data are publicly available at the GEO database and the accession numbers are GSE316292 and GSE317821 (SuperSeries ID: GSE318198).

## ACKNOWLEDGEMENTS

We thank Dr. Frederic Bonnet, director of MDI Biological Laboratory’s Light Microscopy Facility Core (RRID:SCR_019166).

We are also grateful to Comparative Animal Models Core at MDIBL and Lynne Beverly for taking care of housing and feeding of the animals. Also, INBRE summer students at MDIBL, Jacquelin Morin, Mildred J. Prince, Braden Jondro for helping with animal husbandry as well.

We thank Dr. Stephen Sampson, Grant Writer at MDIBL for proofreading this manuscript.

---

**Figure S1.**
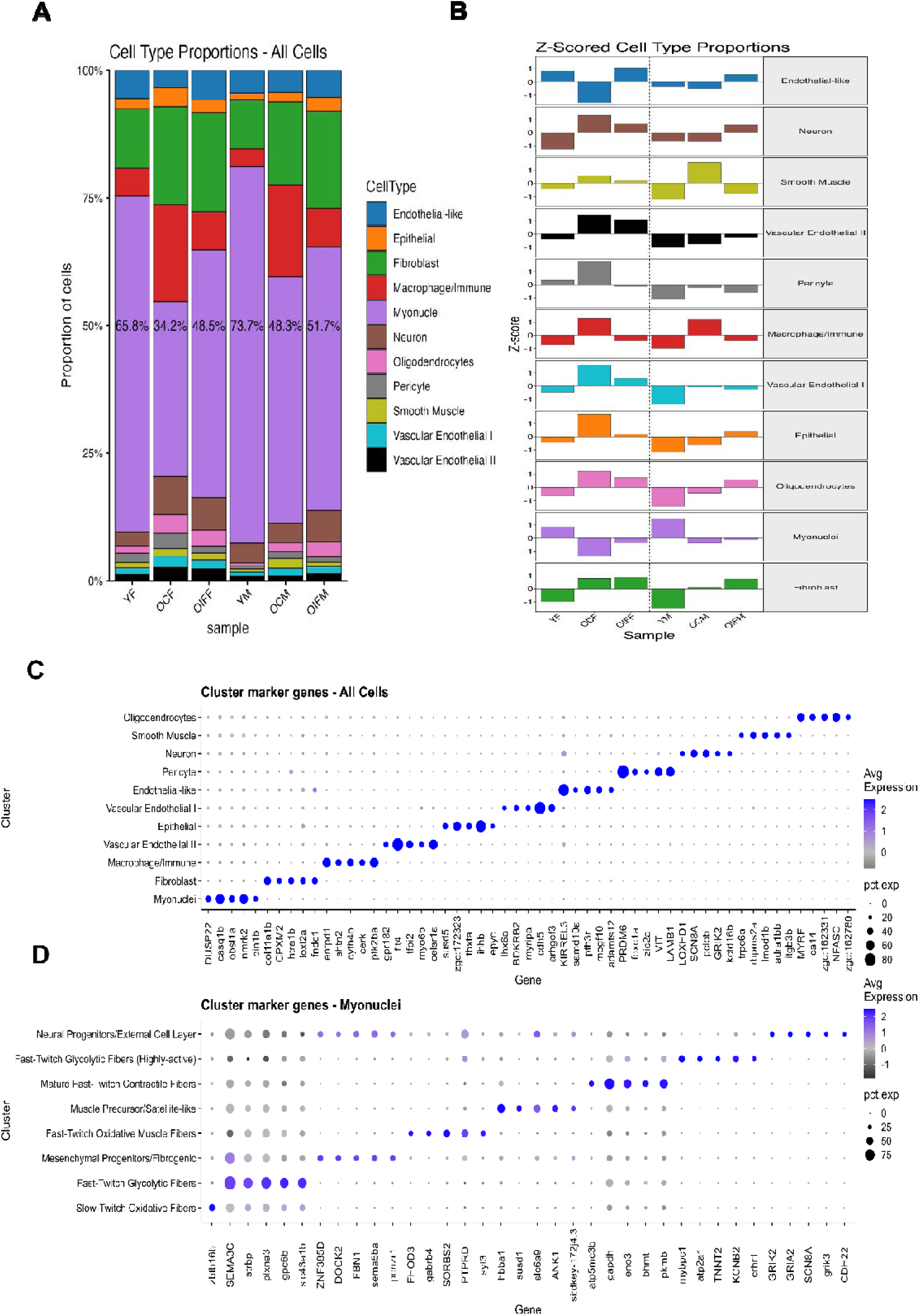
Single-nuclei RNA sequencing of ATK muscle tissue. **A)** Normalized Z-scored comparison of different cell types of muscle tissue between Y, OC, and OIF, male and females. **B)** Stacked bar plots showing proportions of different muscle cell types in Y, OC, OIF, males and females. **C,D)** Dot plots of Seurat-generated markers for different cell types in all cell types, and in the myonuclei subset, respectively.

**Figure S2.**
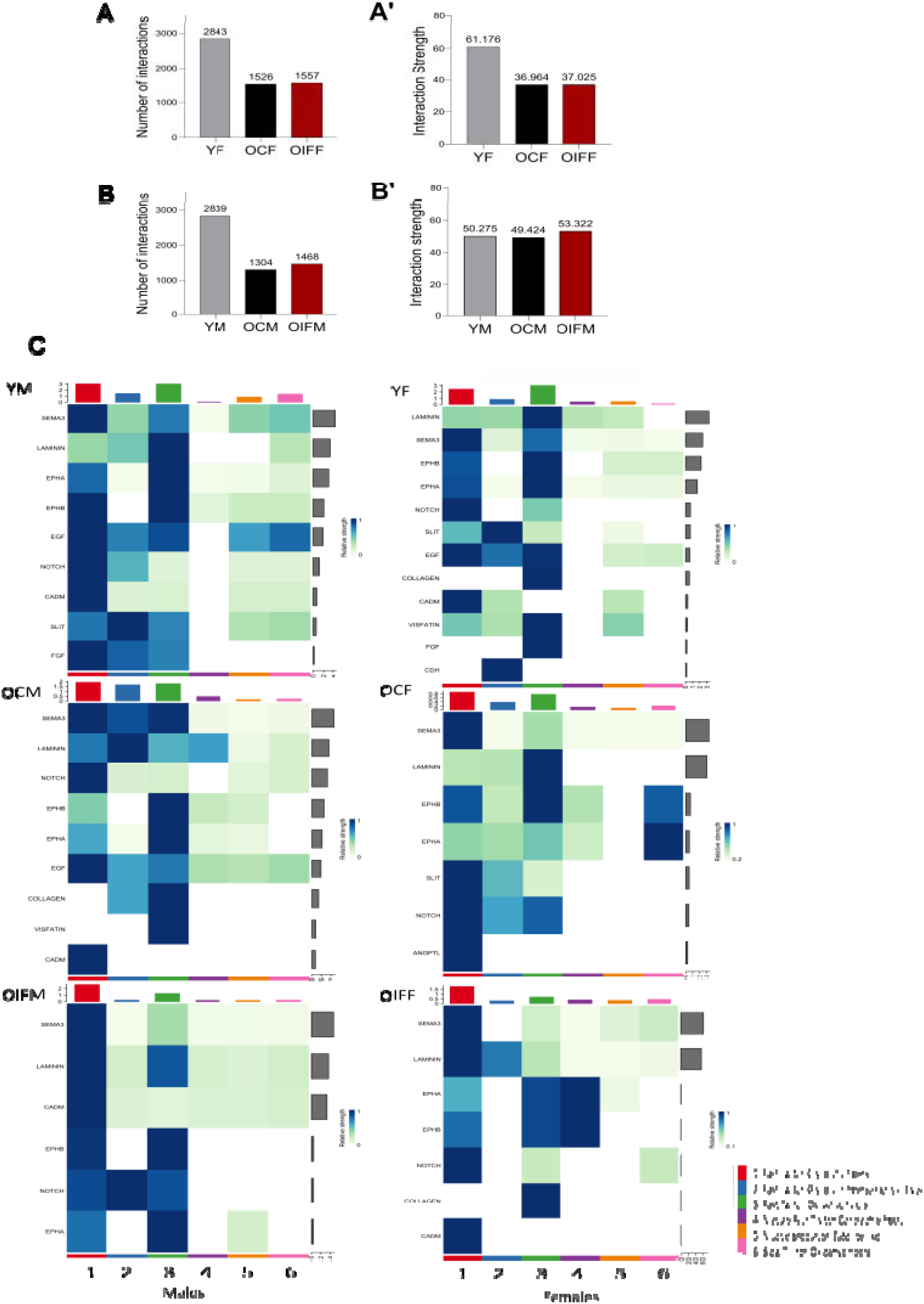
CellChat analysis after snRNA sequencing of ATK muscle tissue. **A, B)** Bar graph with total number of cell-cell interactions across Y, O, and OIF, males and females, respectively. **A’, B’)** Bar graphs with interaction strengths of cell-cell interactions across Y, OC, and OIF, males and females, respectively. **C)** Heat maps of incoming cell-signaling pathways in 6 different myonuclei fiber types across Y, OC, and OIF samples in males (left) and females (right).

**Figure S3.**
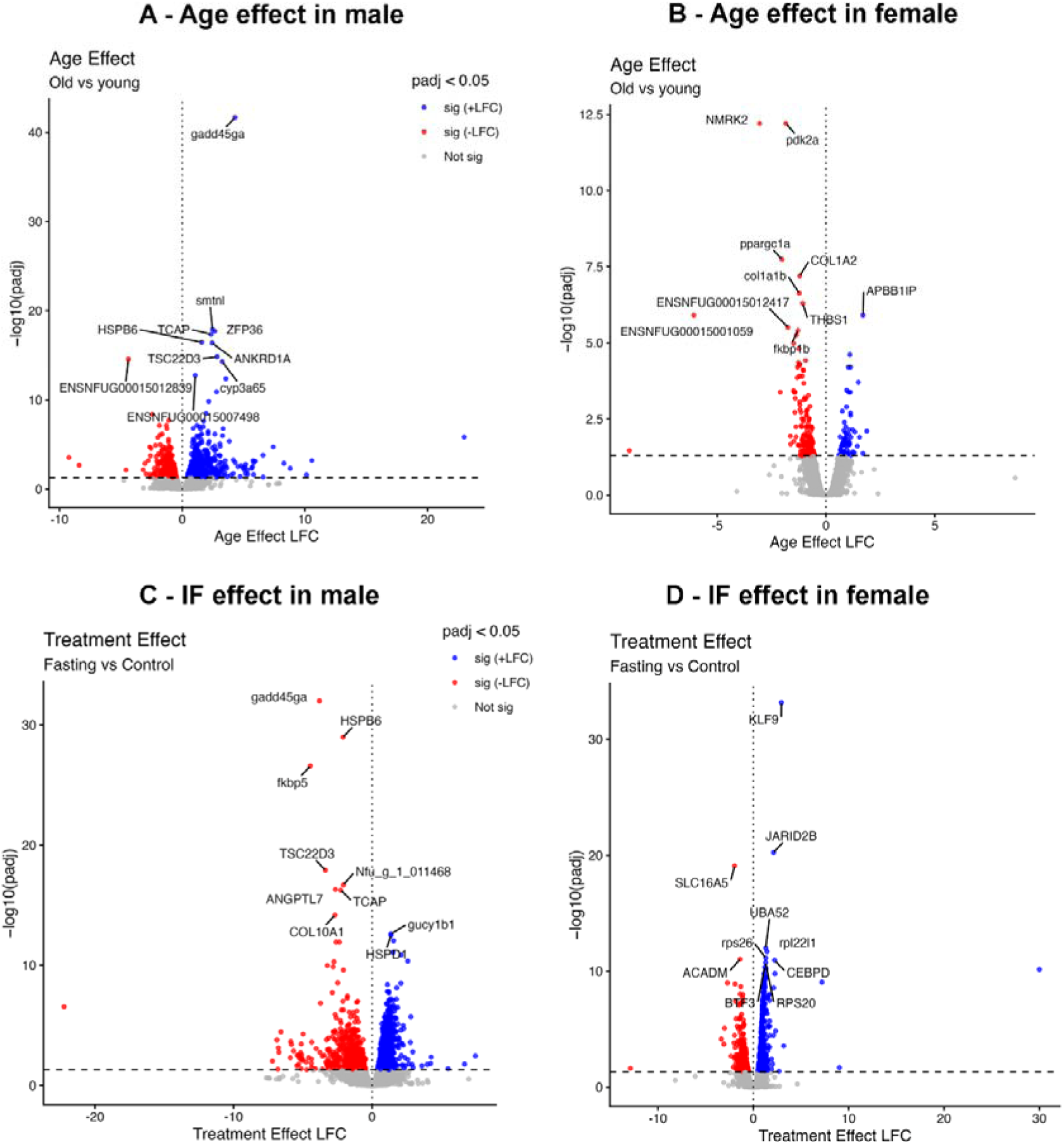
Graphical representation of bulk RNA-seq of ATK muscle tissue. **A, B)** Graphical representation of bulk RNA-seq analysis and variation in gene-expression profiles in young (n=3 males and n=3 females) vs. old (n=3 males and n=3 females) fish, males (A) and females (B), respectively. **C) D)** Graphical representation of bulk RNA-seq analysis and variation in gene-expression profiles in old control (n=3 males and n=3 females) vs. old-after-IF (n=3 males and n=3 females) fish, males (D) and females (E), respectively.

